# Broadband slow-wave modulation in posterior and anterior cortex tracks distinct states of propofol-induced unconsciousness

**DOI:** 10.1101/712604

**Authors:** Emily P. Stephen, Gladia C. Hotan, Eric T. Pierce, P. Grace Harrell, John L. Walsh, Emery N. Brown, Patrick L. Purdon

## Abstract

A controversy 5 has developed in recent years over the role that frontal and posterior cortices play in mediating consciousness and unconsciousness. One hypothesis proposes that posterior sensory and association cortices are the principal mediators of consciousness, citing evidence that strong slow-wave activity over posterior cortex during sleep disrupts the contents of dreaming. A competing hypothesis proposes that frontal-posterior interactions are critical to ignite a conscious percept, since activation of frontal cortex appears necessary for perception and can reverse unconsciousness under anesthesia. In both cases, EEG slow-waves (< 1 Hz) are considered evidence that up- and down-states are disrupting cortical activity necessary for consciousness. Here, we used anesthesia to study the interaction between the slow-wave and higher frequency activity in humans. If slow-waves are derived from underlying up and down-states, then they should modulate activity across a broad range of frequencies. We found that this broadband slow-wave modulation does occur: broadband slow-wave modulation occurs over posterior cortex when subjects initially become unconscious, but later encompasses both frontal and posterior cortex when subjects are more deeply anesthetized and likely unarousable. Based on these results, we argue that unconsciousness under anesthesia comprises several shifts in brain state that disrupt the sensory contents of consciousness distinct from arousal and awareness of those contents.

**Significance Statement:** The roles of frontal and posterior cortices in mediating consciousness and unconsciousness are controversial. Disruption of posterior cortex during sleep appears to suppress the contents of dreaming, yet activation of frontal cortex appears necessary for perception and can reverse unconsciousness under anesthesia. We studied the time course of regional cortical disruption, as mediated by slow-wave modulation of broadband activity, during anesthesia-induced unconsciousness in humans. We found that broadband slow-wave modulation covered posterior cortex when subjects initially became unconscious, but later encompassed both frontal and posterior cortex when subjects were deeply anesthetized and likely unarousable. This suggests that unconsciousness under anesthesia comprises several shifts in brain state that disrupt the contents of consciousness distinct from arousal and awareness of those contents.

## Introduction

A controversy has arisen over the last several years regarding the role that frontal and posterior cortices play in mediating consciousness. The “posterior hot zone” hypothesis proposes that posterior sensory and sensory association cortices are the principal mediators of consciousness, distinct from prefrontal cortex (*1–3*). In contrast, some groups argue that frontal-posterior interactions are critical to ignite a conscious percept (*4–6*). The controversy has recently expanded to include studies of unconsciousness, and how functional impairment in frontal and posterior regions relates to loss of consciousness (*4*, *7*).

Different states of unconsciousness such as anesthesia, NREM sleep, and coma, have distinct electrophysiological signatures. One feature that is common to all of them, however, is slow-wave activity, seen in the electroencephalogram (EEG) as large deflections alternating at approximately 1 Hz. These waves are thought to be large-scale indicators of underlying cortical up- and down-states in which neurons cycle between sustained periods of depolarization (up-states) and hyperpolarization (down-states) (*8*, *9*). In the depolarized up-state, neurons may fire, but in the hyperpolarized down-states, neurons are silent.

While slow-wave activity during unconsciousness is spatially widespread, recent studies have examined how the spatial distribution of slow-wave dynamics relates to states of unconsciousness. During NREM sleep, slow-wave power over posterior areas is stronger during non-dreaming sleep compared to dreaming sleep (*3*). In contrast, frontal areas have been implicated in anesthesia-induced unconsciousness. Pharmacological stimulation of prefrontal cortex can restore behavioral arousal in anesthetized animals, alongside an apparent decrease in slow oscillation power (*10*). Anesthesia-induced slow oscillations modulate frontal alpha oscillations at different phases (“troughmax” and “peakmax” dynamics) depending on the depth of anesthesia (*7*,*11–13*): when frontal alpha power is strongest at the peak of the slow-wave (peakmax), it seems to reflect a deep anesthetic state in which subjects cannot be aroused from unconsciousness even in the presence of painful stimuli (*7*, *12*). These data suggest that different behavioral states sharing similar slow-wave power can be dissociated from each other based on the modulatory influence of the slow-wave on higher frequency rhythms. Could this shift in perspective from slow-wave power to slow-wave modulation be used to reconcile the roles of posterior and frontal cortices in mediating unconsciousness?

While previous cross-frequency modulation analyses during unconsciousness have focused on the influence of the slow-wave on the alpha rhythm, the interpretation of slow-waves as cortical up- and down-states would suggest that EEG power at all frequencies should be coupled to the slow-wave, since up- and down-states affect both rhythmic and non-rhythmic neural activity (*14*). Following this idea, we introduce a new analysis method to track the modulatory influence of the slow-wave across a broad range of frequencies measured in the EEG. We use this method to analyze slow-wave modulation in different cortical regions across different states of propofol-induced unconsciousness. First, we find that frontal peakmax dynamics are a broadband phenomenon, suggesting that peakmax dynamics reflect underlying cortical up- and down-states. Moreover, we find that posterior and frontal regions exhibit this broadband modulation by the slow-wave at different states of unconsciousness: posterior regions at lighter levels of unconsciousness and frontal regions at deeper levels of unconsciousness. This result supports the idea that anesthesia-induced unconsciousness is not a single state but rather multiple states, each showing different levels of posterior and frontal cortical disruption alongside corresponding differences in conscious processing and behavioral arousability.

## Results

We quantified cross-frequency coupling using a correlation between the band-pass filtered slow voltage (0.1-4 Hz) and the instantaneous amplitude of the band-pass filtered high-frequency signal (Figure 1, Panel A). This correlation provides a one-dimensional summary of cross-frequency coupling: if the correlation is positive, it means that the high frequency amplitude is higher when the slow voltage is positive (peakmax coupling); if the correlation is negative, it means that the high frequency amplitude is higher when the slow voltage is negative (troughmax coupling). We can then vary the frequency band used for the high frequency amplitude, to see how different frequencies relate to the slow-wave. Using this approach, we can look at how peakmax and troughmax dynamics vary across electrodes and frequencies during the transition to unconsciousness.

**Figure 1:**
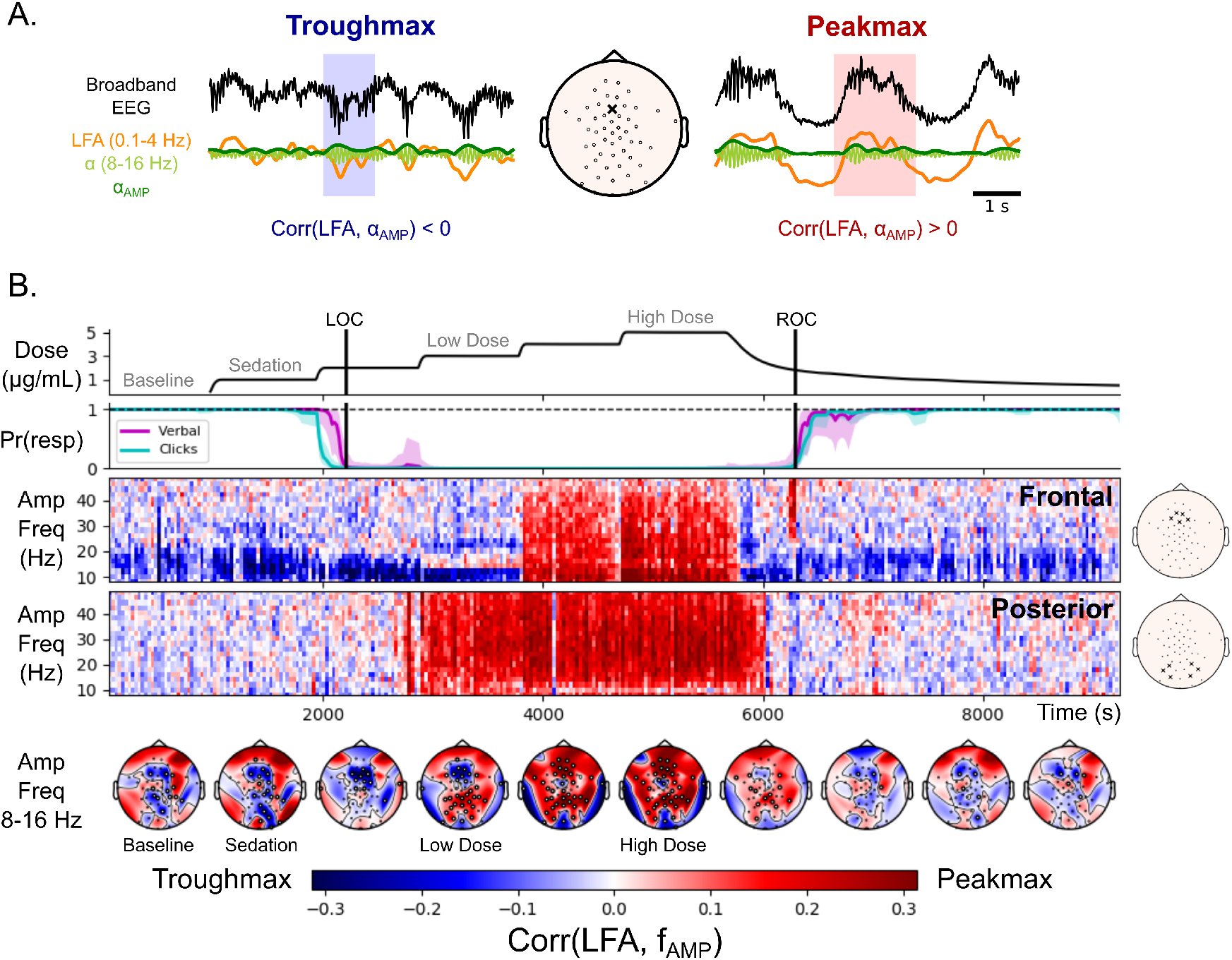
Peakmax coupling is broadband, and it begins after loss of consciousness on posterior electrodes before frontal electrodes. (A) Quantification of cross-frequency coupling. Two example traces from a frontal electrode (middle, “x”): “troughmax” (left) and “peakmax” (right). For each, the broad-band EEG voltage (black) is bandpass filtered into low-frequency activity (LFA, orange) and high frequency activity (*α*, light green). The instantaneous amplitude of the high frequency signal is computed using the analytic signal (*α*_*AMP*_, dark green). The cross-frequency coupling quantified by the correlation between the low frequency activity and the instantaneous amplitude of the high frequency signal. On the left the correlation is negative (troughmax, blue), and on the right the correlation is positive (peak-max, red). (B) Summary of cross-frequency coupling for one subject. Top Panel: the target effect site concentration of propofol administered over the course of the session. Vertical lines indicate Loss of Consciousness (LOC) and Return of Consciousness (ROC). Second Panel: the probability of response curves for the two types of auditory stimuli (Verbal and Clicks). LOC was defined when the probability of response fell below 5% and remained for at least 5 minutes, and ROC was defined when the probability of response went above 5% and remained there for at least 5 minutes. Third and Fourth Panels: the cross-frequency coupling between the LFA and a range of amplitude frequencies (y-axis), for every 30 s interval in the session, for a set of five frontal electrodes (third panel) and a set of six posterior electrodes (fourth panel) with scalp positions indicated on the right. Fifth Panel: Scalp distribution of the cross-frequency coupling between the LFA and the 8-16 Hz amplitude.

Figure 1 (Panel B) shows how this plays out for a representative subject: the subject was administered propofol in increasing doses every 14 minutes while performing an auditory task with a button-click response. When the propofol dose was sufficiently high, the subject stopped responding to the stimulus (LOC). After 5 increasing levels of propofol, the dose was reduced and eventually the subject started responding to the stimulus again (ROC).

The correlation-based cross-frequency coupling over frontal electrodes reaffirms the basic result from previous work on the alpha rhythm: at 10 Hz, coupling to the slow-wave begins as troughmax dynamics, switches to peakmax at higher doses, and transitions back through troughmax during recovery. By looking beyond 10 Hz, however, we can see that the troughmax dynamics begin well before LOC at higher frequencies up to about 20 Hz, frequencies that show higher power during propofol sedation in which subjects remain conscious (*11*). This is also consistent with the observation that patients may be conscious during troughmax dynamics (*7*, *11*, *12*).

Perhaps the most striking result in the cross-frequency coupling is that the peakmax pattern extends across all frequencies (5-50 Hz). Since there are no narrow-band rhythms in the signal other than alpha and slow, this result suggests that non-rhythmic broadband activity is coupled to the slow-wave. This is consistent with an interpretation in which peakmax dynamics reflect underlying cortical up- and down-states, wherein cortical activity, whether rhythmic or not, is shut down during the trough of the slow-wave (*14–16*).

Comparing the cross-frequency coupling between the frontal and posterior electrodes, we can see that a broadband peakmax pattern develops in posterior channels shortly after loss of consciousness, at the same time as frontal troughmax. That is, even before frontal channels transition into peakmax dynamics, posterior channels already show broadband coupling to the slow-wave. In all subjects, the onset of posterior peakmax dynamics occurred after LOC and before the onset of frontal peakmax dynamics (See Figures S2-S10). This result is further shown in the topographic maps of the 8-16 Hz cross-frequency coupling by level: shortly after loss of consciousness, a collection of frontal electrodes still have troughmax coupling while electrodes over posterior electrodes show peakmax coupling. As the propofol level is increased, the frontal electrodes begin to participate in the peakmax coupling. This suggests that broadband silencing of cortical activity by the slow-wave begins first over posterior areas before extending frontally at higher doses of propofol.

For the rest of the analyses, we identified four levels of interest: (1) Baseline, before the administration of propofol; (2) Sedation, the level before the subject stopped responding; (3) Unconscious Low Dose, the first level after the subject stopped responding; and (4) Unconscious High Dose, the highest level of propofol that did not result in burst suppression, a coma-like state beyond what is required for unconsciousness. This allowed us to combine information across subjects, given that the timecourse of LOC and ROC were different for each subject.

Since the patterns of cross-frequency coupling across frequency varied by electrode, propofol level, and subject, we wanted to identify the coupling patterns that best summarized this activity. To do this, we used a non-centered principal component analysis (PCA) to decompose the cross-frequency coupling patterns into principal frequency modes (Figure 2). The principal modes are orthogonal patterns across frequency that capture successively smaller proportions of the total energy in the matrix representing cross frequency coupling by frequency, electrode, propofol level, and subject (Figure S11).

**Figure 2:**
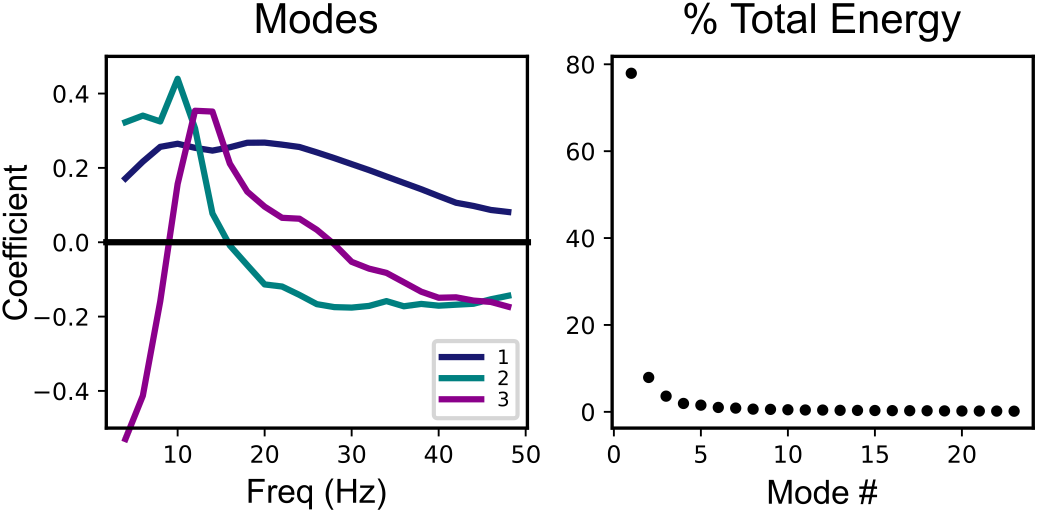
Broadband effects dominate the cross-frequency coupling patterns across subjects, electrodes, and anesthetic drug levels. Here we show the results of a non-centered PCA analysis on the cross-frequency coupling patterns (i.e., to identify frequency-dependent functions describing the modulation of the high frequency signal by the LFA). The analysis was performed over all sensors, subjects, and propofol levels of interest (baseline, sedation, unconscious low dose, and unconscious high dose). Left: The first three principal modes, representing the patterns across frequency that capture the greatest percentage of total energy in the coupling patterns. Right: the percentage of total energy captured by each principal mode, in order from greatest to least. The first principal mode captures 77% of the total energy of the coupling patterns.

The first principal mode is positive for all frequencies, so it represents broadband effects in the cross-frequency coupling, and it captures 78% of the total energy. The second and third principal modes, which have narrow-band peaks in the alpha and beta frequency ranges, respectively, capture much less of the energy. Hence the entrainment of broadband cortical activity to the slow-wave is by far the biggest contributor to the cross-frequency coupling during propofol anesthesia.

This dominant first frequency mode is of particular interest, because it can capture the broadband peakmax pattern observed in the individual subject summaries (Figure 1 and Figures S2-S10). In particular, we would like to use the first principal mode to describe the spatial distribution of the broadband peakmax phenomenon among the cortical generators of the EEG during progressively higher doses of propofol. In order to do so, we performed the cross-frequency coupling analysis on the source-localized EEG signals, and projected the resulting cross-frequency coupling patterns across cortical location onto the first principal mode from the sensor-space analysis described above. Figure 3 shows the resulting projections, morphed to the Freesurfer-average surface (*17*) and averaged across subjects. On these surfaces, a positive projection (red) represents broadband peakmax coupling to the slow-wave.

**Figure 3:**
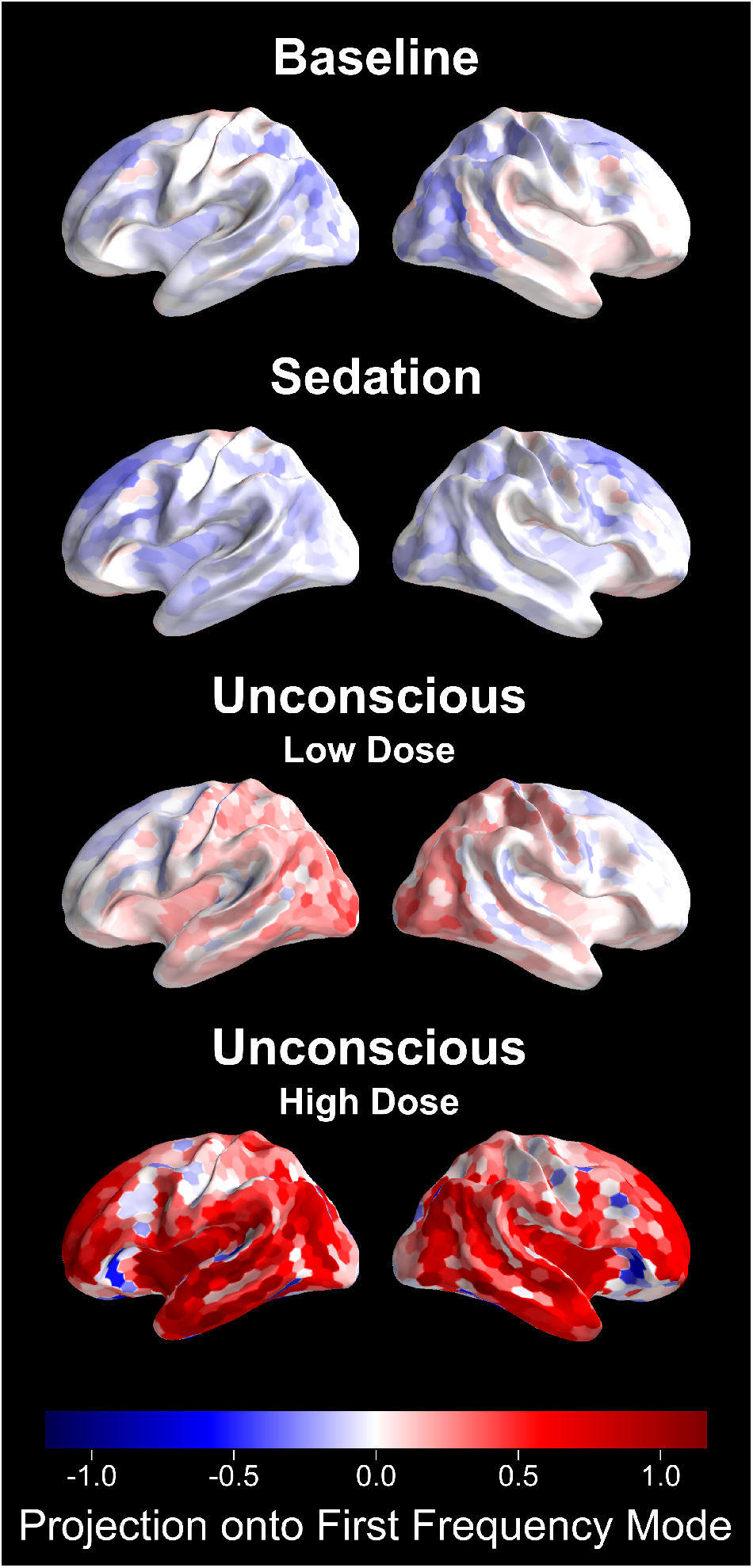
Broadband peakmax occurs first over posterior cortical regions during unconsciousness at lower doses of propofol, and extends to encompass to frontal cortical regions at higher propofol doses. These figures s how the projection of the source-space cross-frequency coupling patterns onto the first principal mode for the four levels of interest. Positive values reflect broadband cross-frequency coupling to the peak of the slow-wave, while negative values represent broadband coupling to the trough of the slow-wave. We present the average coupling across all subjects.

Consistent with the individual subject sensor-space results, broadband peakmax coupling to the slow-wave is present over posterior cortical areas during the Unconscious Low Dose condition. It strengthens and spreads to frontal cortical areas during the Unconscious High Dose condition. In Figure 4, we can see that during the Unconscious Low Dose condition the parietal, temporal, and occipital lobes all have broadband peakmax coupling, but the frontal lobe has virtually no broadband coupling to the slow-wave. During the Unconscious High Dose condition, in contrast, the frontal lobe joins the other lobes in broadband peakmax coupling.

**Figure 4:**
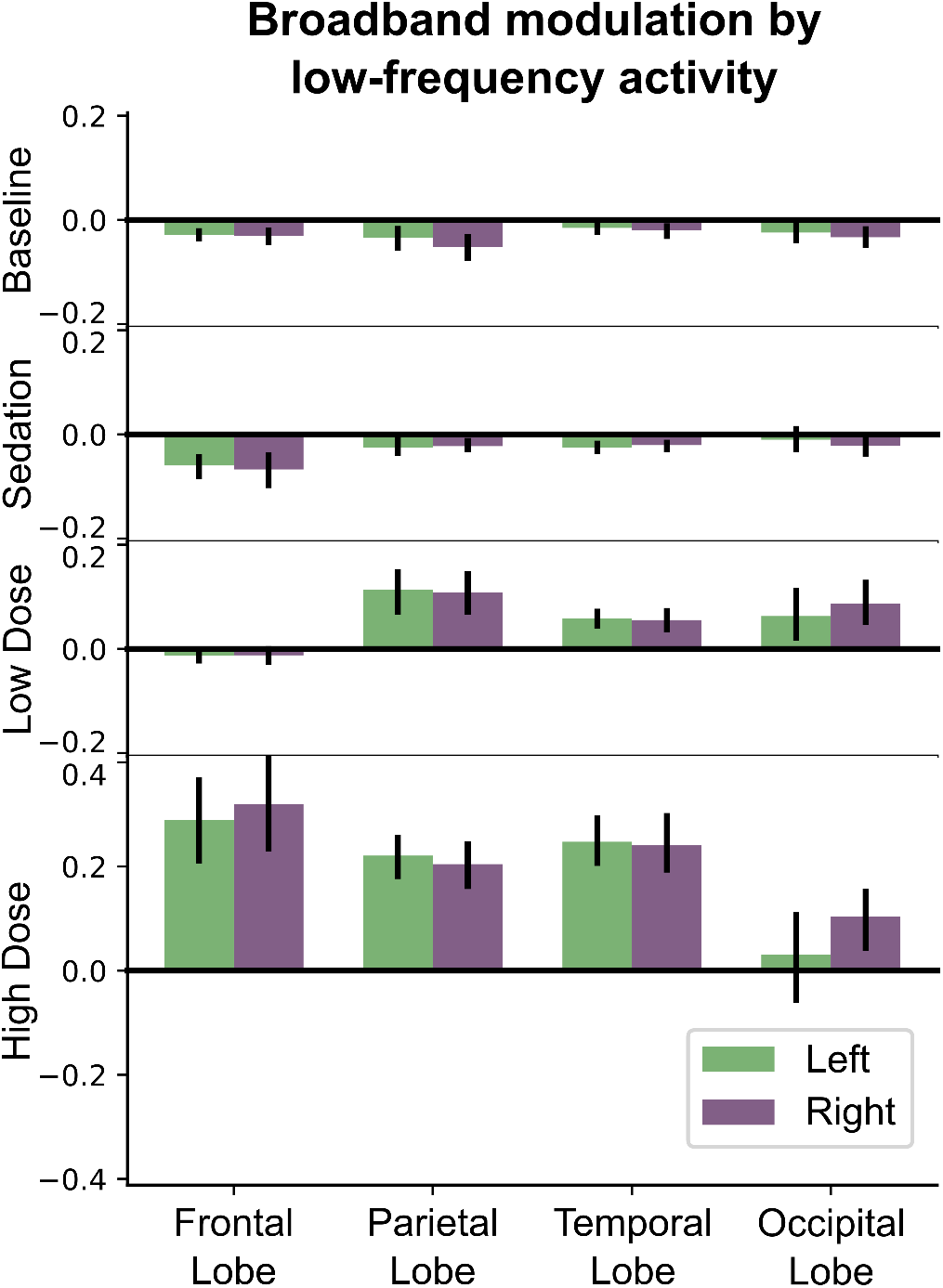
During unconsciousness at low propofol doses, the broadband peakmax effect occurs over the occipital, parietal, and temporal lobes, but not the frontal lobe. Here we show cross-frequency coupling patterns estimated across each cortical lobe using source localization, projected onto the first principal mode and averaged across subjects. Error bars represent 95% confidence intervals, estimated using a bootstrap across subjects.

Together these results suggest that broadband peakmax coupling to the slow-wave is a major contributor to cross-frequency coupling dynamics during propofol anesthesia (Figure 2). Furthermore, posterior cortical areas are the first to exhibit broadband peakmax coupling after loss of consciousness, followed by frontal areas at high doses of propofol (Figures 3 and 4).

## Discussion

We propose that the broadband coupling to the slow-wave that we see in the EEG signal is a macro-scale indicator of regional cortical up- and down-states. Population spiking activity has a broadband effect on the neural power spectrum (*15*, *16*), so the entrainment of population spiking activity to the slow-wave during cortical up- and down-states should cause broadband power to be strongest at a particular phase of the slow-wave. This idea is supported by micro-scale evidence in electrocorticography and local field potential data showing that up-states are associated with simultaneous increases in both population spiking activity and broadband power (<50 Hz) (*14*). While it is common to assume that increased slow power in the EEG always corresponds to up- and down-states in cortical firing, our data suggests that the slow-wave must reach a critical amplitude, which may vary by location, before it entrains the population spiking activity enough to produce broadband modulation. Posterior areas cross this threshold at lower doses of propofol than frontal areas. The idea of a critical threshold for the slow-wave is consistent with the slow-wave activity saturation hypothesis (*18*), in which saturation of the slow-wave power accompanies a disconnection between thalamus and sensory cortex. We hypothesize that the cortical up- and down-states themselves provide a mechanism for this sensory isolation. Under this interpretation, broadband peakmax dynamics signify that local cortical activity is being disrupted by up- and down-states on a scale large enough to be detected in the EEG.

The ability to monitor a patient’s state of unconsciousness during general anesthesia has obvious practical benefits, but identifying principled methods for doing so remains an open question. We can make a distinction between unconscious states in which patients can be aroused to consciousness by external stimuli (arousable), versus those in which patients cannot be aroused (unarousable). Some recent data suggest indirectly that patients can be arousable when frontal alpha rhythms are strongest at the the trough of the slow-wave (troughmax), but are unarousable when the frontal alpha rhythms are strongest at the peak of the slow-wave (peakmax) (*7*,*11*,*12*). Here we show that when frontal alpha rhythms are coupled to the peak of the slow-wave (peak-max), broadband spectral power over both frontal and posterior cortices, is also coupled to the slow-wave. Hence unarousability appears to be associated with broadband coupling to the slow-wave over both frontal and posterior cortex. At lower anesthetic doses in which patients are unconscious but potentially arousable, we find that the posterior cortex engages in broad-band peakmax dynamics while the prefrontal cortex participates in (alpha-band) troughmax dynamics. Thus, these different spatial patterns of cross-frequency coupling appear to indicate different states of unconsciousness. Knowledge of these states could be used by anesthesiologists to establish different arousable or unarousable states of unconsciousness appropriate to the clinical circumstances.

The “posterior hot zone” (*1–3*) and fronto-posterior (*4–6*) hypotheses propose starkly different roles for posterior and frontal circuits in mediating consciousness. Anesthesia introduces another dimension to this problem in that unconscious patients may or may not be arousable by external stimuli depending on the drug dose. Along this dimension, the prefrontal cortex appears to play a central role, in which stimulation of prefrontal cortex can jump-start consciousness (*10*). Our results would seem to harmonize these viewpoints, in that the disruption of frontal versus posterior cortices coincides with different states of unconsciousness. First, both the posterior hot zone and fronto-posterior hypotheses predict that disruption of posterior areas should degrade the contents of consciousness. In our results, posterior areas are the first to show broadband peakmax at low doses of propofol, when subjects are unconscious but may be arousable. These posterior areas appear strikingly similar to those with higher slow-wave power during non-dreaming sleep (*3*). This is consistent with the interpretation that posterior cortical regions mediate the contents of consciousness (*2*), which are disrupted by alternating up- and down-states during unconsciousness (*3*, *14*). Second, frontal areas develop broadband peakmax coupling at higher doses of propofol, in a state associated with unarousability. This is consistent with the interpretation that prefrontal cortex is required for behavioral arousal to consciousness (*10*) and that alternating up- and down-states in prefrontal cortex prevent this arousal.

The main result of this study is that brain-wide cross-frequency coupling analysis captures reliable but spatially-varying changes in brain dynamics during the transition to unconsciousness and unarousability under propofol anesthesia. We interpret broadband coupling to the peak of the slow-wave to be a macro-scale indicator of up- and down-states in the micro-scale cortical activity that serve to disrupt cortical function. This functional disruption encompasses both frontal and posterior cortical areas at different states of unconsciousness. Our results suggest that unconsciousness under anesthesia is not a singular phenomenon but rather involves several distinct shifts in brain state that disrupt posterior cortical processing of the “contents” of consciousness distinct from prefrontal arousal and awareness of those contents.

## Online Methods

### Experimental Design

This study analyzes data that has been previously described in (*11*) and (*13*). All procedures were approved by the Human Research Committee at the Massachusetts General Hospital.

Ten healthy volunteers between the ages of 18 and 36 participated. Prior to the study, structural MRI scans were obtained for each subject (Siemens Trio 3 Tesla, T1-weighted magnetization-prepared rapid gradient echo, 1.3-mm slice thickness, 1.3 1 mm in-plane resolution, TR/TE = 2530/3.3 ms, 7 flip angle). Cortical reconstruction was performed with the FreeSurfer image analysis suite (*17*, *surfer.nmr.mgh.harvard.edu*).

After at least 14 minutes of eyes-closed rest, subjects were administered propofol anesthesia, increasing propofol dosage every 14 minutes to target 1, 2, 3, 4, or 5 μgmL^−1^ effect-site concentration. During emergence, propofol dose was decreased in three 14-minute steps targeting effect-site concentrations of 0.5 μgmL^−1^, 1.0 μgmL^−1^, and 1.5 μgmL^−1^ less than the dose at which the subject stopped responding to the auditory cues. Electroencephalogram (EEG) recordings were collected with a high density (64 channel) BrainVision MRI Plus system (Brain Products) with a sampling rate of 5,000 samples per second and the electrode positions were digitized (Polhemus FASTRACK 3D).

Throughout the study, subjects performed a task in which they responded to an auditory click or word with a finger-press of a button. Loss of consciousness (LOC) was defined when a state-space model estimated a less than 5% probability of a correct response and remained so for at least 5 minutes (*11*). Similarly, return of consciousness (ROC) was defined as the first time during emergence when the state-space model reached a 5% probability of a correct response and remained so for at least 5 minutes.

### EEG preprocessing

Channels with severe, persistent artifacts were identified by visual inspection and removed from the analysis. We also removed electrodes that showed signs of being electrically bridged according to the eBridge software package (*19*).

The cross-frequency coupling analysis described below uses versions of the signal that have been bandpass filtered in the slow band (0.1-4 Hz) and a set of higher bands between 4 and 50 Hz, each 2 Hz in bandwidth with a 1 Hz transition band. We filtered the data for the entire session into these bands using a zero-phase FIR filter and downsampled to 200 samples per second. We chose the upper limit of the higher bands, 50 Hz, in order to avoid line noise and capture the frequencies with the largest EEG power (*20*).

For sensor-level analyses, we applied a Laplacian reference based on first nearest neighbors after filtering (*21*).

Because of the stepped structure of the propofol dosage, subjects had approximately 12 minutes at each dosage with constant predicted effect-site concentration. We call these periods “levels”, and defined four levels of interest: Baseline, Sedation, Unconscious Low Dose, and Unconscious High Dose. Baseline was defined as the period before the onset of propofol, Sedation was defined as the level prior to LOC, Unconscious Low Dose was defined as the first full level after LOC, and Unconscious High Dose was defined as the highest level of propofol or the level before burst suppression (for subjects who entered burst suppression). For analyses performed on these levels, we selected 10 artifact-free epochs 30 s long from each level for each subject. Two of the subjects stopped responding during the first level of propofol, so they did not have a sedation period. As a result, figures summarizing across subjects represent 10 subjects for Baseline, Low Dose, and High Dose, and 8 subjects for Sedation.

### Cross-frequency coupling analysis

Under propofol, high frequency amplitude is much more likely to couple to the peak or the trough of the slow-wave than to the rising or falling phases (*11*) (Figure S1). Hence we summarize cross-frequency coupling using a metric that is either positive (peakmax: high frequency amplitude is higher during the peak of the slow-wave) or negative (troughmax: high frequency amplitude is higher during the trough of the slow-wave). In particular, we use the Pearson correlation coefficient between the instantaneous amplitude of the band-passed high frequency signal and the band-passed slow voltage (0.1-4 Hz) (*22*):

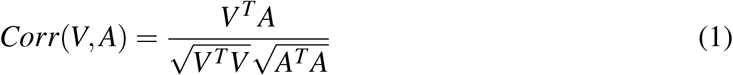

where *V* is a column vector representing a timeseries of the slow voltage and *A* is a vector representing the corresponding timeseries of the instantaneous amplitude of the high frequency band. The instantaneous amplitude was computed using the magnitude of the analytic signal of the bandpass filtered signal. Before computing the correlation, the instantaneous amplitude for each 30 s epoch was centered by subtracting the mean. *V* was not explicitly centered because its expected value is zero.

In order to find the average correlation across space (electrodes, source locations) and/or time (multiple 30 s epochs), the dimensions to be averaged are stacked vertically in the *V* and *A* column vectors in Equation 1. Since the correlation represents the covariance between the signals (in the numerator, *V*^*T*^*A*) normalized by the standard deviations of each signal (in the denominator, 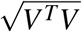 and 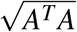), stacking the signals vertically has the effect of estimating the covariance and standard deviations separately before performing the normalization.

Note that this analysis does not involve estimating the phase of the slow-wave, a typical step in phase-amplitude coupling analysis (*22*). We chose not to estimate phase for three reasons. First, phase-amplitude coupling in propofol clusters around zero (slow-wave peak, when the signal is positive) and *π* (slow-wave trough, where the signal is negative) (*11*) (see also Figure S1). Second, the slow-wave is thought to reflect up states and down states, regardless of whether it is well-described by a sinusoid. Early in propofol-induced unconsciousness, the slow-wave is often quite irregular in waveform shape and frequency. We chose to use a wider band for the slow-wave, 0.1-4 Hz, in order to capture more of the non-sinusoidal features of the waveform. Under propofol anesthesia, there is no narrow-band delta frequency rhythmic activity (1-4 Hz), so the contribution of this frequency range to the band-passed signal was non-rhythmic, affecting the waveform shape. By using correlation to describe the relationship of the high frequency amplitude to the non-sinusoidal low frequency waveform, we are therefore better able to capture the relationship between high frequency activity and the up and down fluctuations that are apparent in the broadband signal (Figure 1, Panel A). Finally, reducing the problem to one dimension (positive and negative coupling) simplified the analysis of a single location and frequency, allowing us to expand the analysis to other frequencies and locations.

Panel A in Figure 1 shows the computation for a frontal electrode, computing the correlation between the slow voltage (low-frequency activity, LFA) and the alpha band (*α*, 9-11 Hz) during two time windows. For the time window on the left, the alpha amplitude is higher when the slow voltage is negative, leading to a negative correlation (troughmax). For the time window on the right, the alpha amplitude is higher when the slow voltage is positive, leading to a positive correlation (peakmax). We can perform this analysis for all 30 s epochs in the session and for a range of amplitude frequencies (2 Hz bands between 4 and 50 Hz): this results in a modulogram showing the cross-frequency coupling between each frequency and the slow-wave across time in the session for the selected electrode. Similarly, we can compute the correlations for each electrode within propofol levels, and show the spatial distribution of cross-frequency coupling between the alpha band and the slow-wave as a function of propofol level.

We performed the cross-frequency coupling analysis across amplitude bands, time, electrodes, and subjects. We characterize time in two ways. The first was a session-based analysis, in which the coupling was assessed for every 30 s epoch in the session separately, resulting in modulograms such as the frontal and posterior summaries in Panel B of Figure 1: these modulograms represent the average correlations across several electrodes, indicated in the scalp maps to the right. The second way to characterize time was based on propofol level, in which the coupling was assessed over 10 artifact-free 30 s epochs at each propofol level. Hence, each coupling value represents the correlation over 300 seconds of data, and we have one such value for each electrode, level, and subject. This level-based analysis was used for the topoplots in Panel B of Figure 1 and for the Principal Component Analysis described below.

### Source localization

Source localization was performed by minimum norm estimation (*23*) using the MNE-python toolbox (*24*, www.martinos.org/mne/). The noise covariance was assumed to be diagonal, estimated from the EEG using the spectral power above 65 Hz. The source space was an ico-3 decimation of the Freesurfer reconstruction of the cortical surface and the sources were constrained to be perpendicular to the cortical surface. The forward model was computed using a 3-layer boundary element model, eliminating any sources within 5mm of the inner skull surface.

To perform the cross-frequency coupling analysis in source space, the band-passed signals in the slow band and each 2 Hz band between 4 and 50 Hz were source localized using the methods described above. For the cortical surface representations (Figure 3), the cross-frequency coupling analysis described above was applied to each source location for each subject, for the four levels of interest. The average cross-frequency coupling within lobes (Figure 4) used the same technique, but the signals for all of the sources in a lobe were combined: the 10 epochs (for each level) for all of the sources in the lobe were stacked vertically in the *V* and *A* column vectors in Equation 1.

### Non-centered Principal Component Analysis and Projection into Source Space

To identify patterns of coupling across frequency that explained most of the observed features, we performed a non-centered principal component analysis (*25*) on the sensor-space coupling patterns across propofol levels and subjects. The level-based coupling results for the four levels of interest (baseline, sedation, unconscious low dose, and unconscious high dose) were combined into an aggregate matrix *A*(*f,i*) where *f* indexes the amplitude band and *i* indexes the propofol level, electrode, and subject (i.e. the rows contain all of the levels, electrodes, and subjects stacked). Figure S11 shows the cross-frequency coupling results for two electrodes during the four levels of interest, for all subjects: the goal of the analysis was to decompose these patterns into components that capture their major features across frequencies. Principal component analysis involves a singular value decomposition:

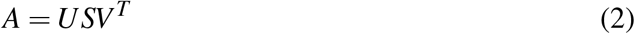

In the decomposition, *U* is a matrix whose columns are the principal modes. The first column is the first principal mode, meaning the pattern across frequencies that captures the most energy in the *A* matrix. The second column is the second principal mode, which captures the most energy in the data after the first mode has been removed. The principal modes are orthogonal, and they all have unit length. In order to preserve the mapping of positive values to peakmax and negative values to troughmax, the principal modes were flipped (multiplied by −1) as necessary so that the largest element would be positive. The *S* matrix is diagonal, and can be used to estimate the percent of the total energy that is explained by the *j*^*th*^ mode:

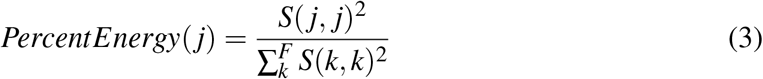

Note that we chose to use non-centered PCA, which decomposes the total energy, rather than centered PCA, which decomposes variance. Non-centered PCA is particularly useful in situations where the origin has special significance that would be lost if the data were centered by subtracting the mean (*25*). For patterns of cross frequency coupling, the origin represents a situation in which the EEG signal contains no coupling to the slow-wave at any frequency. In contrast, the mean represents the average coupling to the slow-wave over subjects, propofol levels, and electrodes (as a function of amplitude frequency). We feel that the origin has more significance than the mean, and as a result the non-centered PCA is more interpretable than the more common centered PCA.

To quantify how the principal modes were represented in source space, we projected the source-space patterns for each subject onto the first sensor-space principal mode. First, the coupling results from the source-space analysis were combined into an aggregate *A*^*source*^(*f,i*) matrix, where *i* indexes the propofol level, source location (or lobe), and subject. Then the projection of *A*^*source*^ onto *U* is:

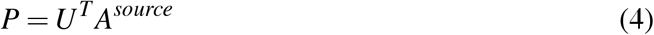

The first row of the resulting *P* matrix contains the projection of source-space coupling patterns (for each propofol level, source location, and subject) onto the first principal mode. Since the first principal mode is relatively constant across frequencies (see Figure 2), the projections onto this mode represent cross-frequency coupling that is broadband. Also, since the first principal mode is positive for all frequencies, positive projections reflect broadband peakmax coupling (all frequencies coupled to the peak of the slow-wave), and negative projections reflect broad-band troughmax coupling (all frequencies coupled to the trough of the slow-wave).

The projections of the (*subject* × *level* × *lobe*) coupling patterns onto the first principal mode were averaged across subjects to yield the estimates shown in Figure 4, representing the average contribution of the first mode to the cross-frequency coupling in that lobe. The 95% confidence intervals were obtained using a bootstrap over subjects. Individual subject results are shown in Figure S12.

To summarize the results across subjects on the entire cortical surface, we morphed the projections of the (*subject* × *level* × *sourcelocation*) coupling patterns onto the first principal mode to the Freesurfer-average surface (*17*) using MNE-python (*24*), and we averaged the resulting maps across subjects. The resulting maps estimate the contribution of the chosen mode to the cross frequency coupling at each cortical location (see Figure 3).

**Supplemental Figure S1: Mean Vector Analysis** To quantify the extent to which the high frequency amplitude couples to the peak or the trough of the slow-wave rather than to the rising or falling phases, we used mean vector analysis (*22*). As described above, 10 30-second artifact-free intervals were chosen in each propofol level for each subject, resulting in 100 intervals spread out over the session for each subject. For each interval, the mean vector estimate of the phase-amplitude coupling is defined as:

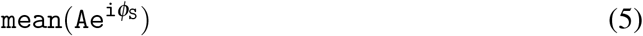

where *A* represents the 30 second timeseries of the instantaneous amplitude of the high frequency band (here, 8-16 Hz) and *ϕ*_*S*_ represents the 30 second timeseries of the instantaneous phase of the low frequency band (here, 0.1-4 Hz). The resulting estimate is a complex number representing both the magnitude of the phase-amplitude coupling and its preferred phase. Figure S1 shows the mean vectors for all 30 second intervals, for all subjects and all electrodes. The results show that the amplitude of the high frequency signals tends to couple to either the peak or trough of the slow oscillation, and helps justify the use of the correlation between slow phase and fast amplitude as a means of quantifying phase-amplitude coupling.

## Acknowledgements

The authors would like to thank Nancy Kopell and Austin Soplata for valuable feedback about theoretical mechanisms, Matti Hämäläinen and Sheraz Khan for helpful discussions on source localization, and Eric Denovellis for help with figure design.

## Author Contributions

Conceptualization, E.P.S., E.N.B., and P.L.P.; Methodology, E.P.S. and P.L.P; Software, E.P.S. and G.C.H.; Formal Analysis, E.P.S. and P.L.P.; Investigation, E.T.P., P.G.H., J.L.W., E.N.B., and P.L.P.; Writing – Original Draft, E.P.S., P.L.P.; Writing – Review & Editing, E.P.S., P.L.P.; Funding Acquisition, E.N.B and P.L.P.

## Data materials and availability

Data and code will be made available in a public database prior to publication.

## Supplementary materials

Materials and Methods Figs. S1 to S13

## References

1. M. Boly, M. Massimini, N. Tsuchiya, B. R. Postle, C. Koch, and G. Tononi, “Are the Neural Correlates of Consciousness in the Front or in the Back of the Cerebral Cortex? Clinical and Neuroimaging Evidence,” The Journal of Neuroscience, vol. 37, pp. 9603–9613, Oct. 2017.

2. C. Koch, M. Massimini, M. Boly, and G. Tononi, “Neural correlates of consciousness: progress and problems,” Nature Reviews Neuroscience, vol. 17, pp. 307–321, May 2016.

3. F. Siclari, B. Baird, L. Perogamvros, G. Bernardi, J. J. LaRocque, B. Riedner, M. Boly, B. R. Postle, and G. Tononi, “The neural correlates of dreaming,” Nature Neuroscience, pp. 1–10, Apr. 2017.

4. G. A. Mashour, “The controversial correlates of consciousness,” Science, vol. 360, pp. 493–494, May 2018.

5. B. Odegaard, R. T. Knight, and H. Lau, “Should a Few Null Findings Falsify Prefrontal Theories of Conscious Perception?,” Journal of Neuroscience, vol. 37, pp. 9593–9602, Oct. 2017.

6. B. v. Vugt, B. Dagnino, D. Vartak, H. Safaai, S. Panzeri, S. Dehaene, and P. R. Roelfsema, “The threshold for conscious report: Signal loss and response bias in visual and frontal cortex,” Science, vol. 360, pp. 537–542, May 2018.

7. A. L. Gaskell, D. F. Hight, J. Winders, G. Tran, A. Defresne, V. Bonhomme, A. Raz, J. W. Sleigh, and R. D. Sanders, “Frontal alpha-delta EEG does not preclude volitional response during anaesthesia: prospective cohort study of the isolated forearm technique,” British Journal of Anaesthesia, Aug. 2017.

8. P. Achermann and A. Borbly, “Low-frequency (<1hz) oscillations in the human sleep electroencephalogram,” Neuroscience, vol. 81, pp. 213–222, Aug. 1997.

9. M. Steriade, A. Nunez, and F. Amzica, “A novel slow (<1 Hz) oscillation of neocortical neurons in vivo: depolarizing and hyperpolarizing components,” The Journal of Neuroscience, vol. 13, p. 3252, Aug. 1993.

10. D. Pal, J. G. Dean, T. Liu, D. Li, C. J. Watson, A. G. Hudetz, and G. A. Mashour, “Differential role of prefrontal and parietal cortices in controlling level of consciousness,” Current Biology, vol. 28, no. 13, pp. 2145–2152, 2018.

11. P. L. Purdon, E. T. Pierce, E. A. Mukamel, M. J. Prerau, J. L. Walsh, K. F. K. Wong, A. F. Salazar-Gomez, P. G. Harrell, A. L. Sampson, A. Cimenser, S. Ching, N. J. Kopell, C. Tavares-Stoeckel, K. Habeeb, R. Merhar, and E. N. Brown, “Electroencephalogram signatures of loss and recovery of consciousness from propofol.,” Proceedings of the National Academy of Sciences of the United States of America, vol. 110, pp. E1142–51, Mar. 2013.

12. E. Brown, P. Purdon, O. Akeju, and J. An, “Using EEG markers to make inferences about anaesthetic-induced altered states of arousal,” British journal of anaesthesia, vol. 121, no. 1, pp. 325–327, 2018.

13. E. A. Mukamel, E. Pirondini, B. Babadi, K. F. K. Wong, E. T. Pierce, P. G. Harrell, J. L. Walsh, A. F. Salazar-Gomez, S. S. Cash, E. N. Eskandar, V. S. Weiner, E. N. Brown, and P. L. Purdon, “A Transition in Brain State during Propofol-Induced Unconsciousness,” Journal of Neuroscience, vol. 34, pp. 839–845, Jan. 2014.

14. L. D. Lewis, V. S. Weiner, E. A. Mukamel, J. A. Donoghue, E. N. Eskandar, J. R. Madsen, W. S. Anderson, L. R. Hochberg, S. S. Cash, E. N. Brown, and P. L. Purdon, “Rapid fragmentation of neuronal networks at the onset of propofol-induced unconsciousness.,” Proceedings of the National Academy of Sciences of the United States of America, vol. 109, pp. E3377–86, Nov. 2012.

15. J. R. Manning, J. Jacobs, I. Fried, and M. J. Kahana, “Broadband Shifts in Local Field Potential Power Spectra Are Correlated with Single-Neuron Spiking in Humans,” Journal of Neuroscience, vol. 29, pp. 13613–13620, Oct. 2009.

16. K. J. Miller, “Broadband Spectral Change: Evidence for a Macroscale Correlate of Population Firing Rate?,” Journal of Neuroscience, vol. 30, pp. 6477–6479, May 2010.

17. A. M. Dale, B. Fischl, and M. I. Sereno, “Cortical Surface-Based Analysis,” NeuroImage, vol. 9, no. 2, pp. 179–194, 1999.

18. R. Ni Mhuircheartaigh, C. Warnaby, R. Rogers, S. Jbabdi, and I. Tracey, “Slow-Wave Activity Saturation and Thalamocortical Isolation During Propofol Anesthesia in Humans,” Science Translational Medicine, vol. 5, pp. 208ra148–208ra148, Oct. 2013.

19. D. M. Alschuler, C. E. Tenke, G. E. Bruder, and J. Kayser, “Identifying electrode bridging from electrical distance distributions: A survey of publicly-available EEG data using a new method,” Clinical neurophysiology: official journal of the International Federation of Clinical Neurophysiology, vol. 125, pp. 484–490, Mar. 2014.

20. P. L. Nunez, R. Srinivasan, and others, Electric fields of the brain: the neurophysics of EEG. Oxford University Press, USA, 2006.

21. A. Cimenser, P. L. Purdon, E. T. Pierce, J. L. Walsh, A. F. Salazar-Gomez, P. G. Harrell, C. Tavares-Stoeckel, K. Habeeb, and E. N. Brown, “Tracking brain states under general anesthesia by using global coherence analysis.,” Proceedings of the National Academy of Sciences of the United States of America, vol. 108, pp. 8832–8837, May 2011.

22. A. B. L. Tort, R. Komorowski, H. Eichenbaum, and N. Kopell, “Measuring phase-amplitude coupling between neuronal oscillations of different frequencies.,” Journal of Neurophysiology, vol. 104, pp. 1195–1210, Aug. 2010.

23. M. Hmlinen, R. Hari, R. J. Ilmoniemi, J. Knuutila, and O. V. Lounasmaa, “Magnetoen-cephalographytheory, instrumentation, and applications to noninvasive studies of the working human brain,” Reviews of modern Physics, vol. 65, pp. 413–497, Apr. 1993.

24. A. Gramfort, “MEG and EEG data analysis with MNE-Python,” Frontiers in Neuroscience, vol. 7, 2013.

25. I. T. Jolliffe, Principal component analysis. Springer, 2002.

