## Supplementary Materials for "Broadband slow-wave modulation in posterior and anterior cortex tracks distinct states of propofol-induced unconsciousness"

**This file includes:**

Figs. S1 to S13

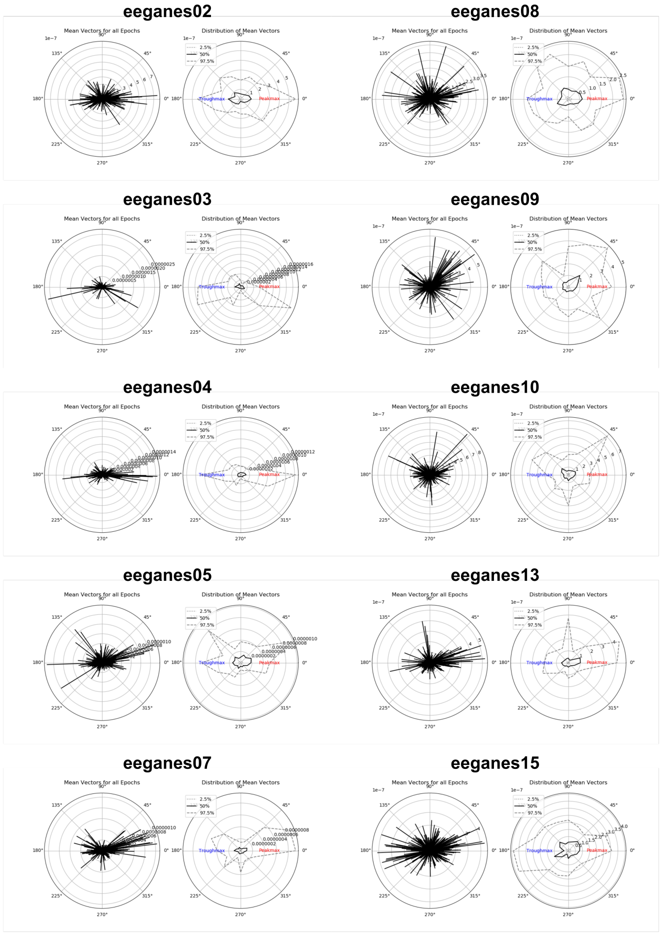


**Fig. S1:** Phase-amplitude coupling between the slow phase (0.1-4 Hz) and the alpha amplitude (8-16 Hz) using mean vector analysis for each subject. For each subject, the polar plot on the left shows the mean vectors for all electrodes for 100 30-second intervals spread over the entire session. The polar plot on the right shows the 2.5%, 50% (median), and 97.5% percentiles of the mean vectors, computed for 16 equally-spaced phase bins. Unlike the correlation-based cross-frequency coupling metric used in the rest of the paper, the mean vector estimate of phase is sensitive to coupling at 90 degrees and 270 degrees, corresponding to the rising phase and the falling phase of the slow oscillation. The results show that the amplitude of the high frequency signals tends to couple to either the peak or trough of the slow oscillation, and helps justify the use of the correlation between slow phase and fast amplitude as a means of quantifying phase-amplitude coupling.

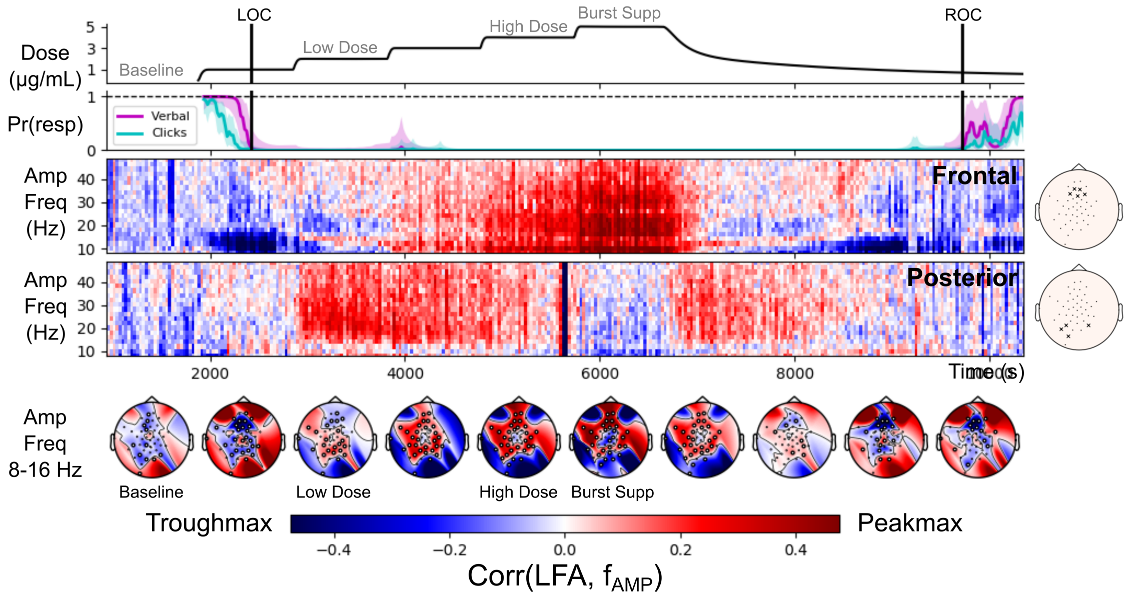


**Fig. S2:** Sensor-level summary for Subject 2. Layout as in Figure 1. Note that this subject entered burst suppression (Burst Supp) at the highest dose of propofol, so the previous level was used for the Unconscious High Dose condition.

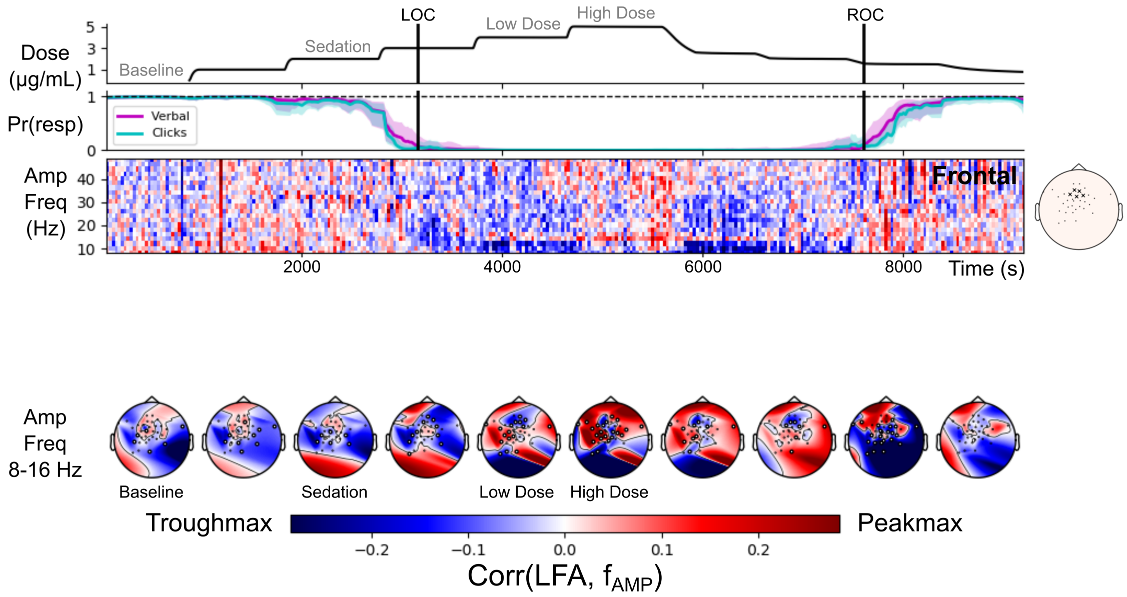


**Fig. S3:** Sensor-level summary for Subject 3. Layout as in Figure 1. A large number of posterior electrodes were excluded for this subject due to electrical bridging (seee Methods), so the posterior cross-frequency coupling has been omitted.

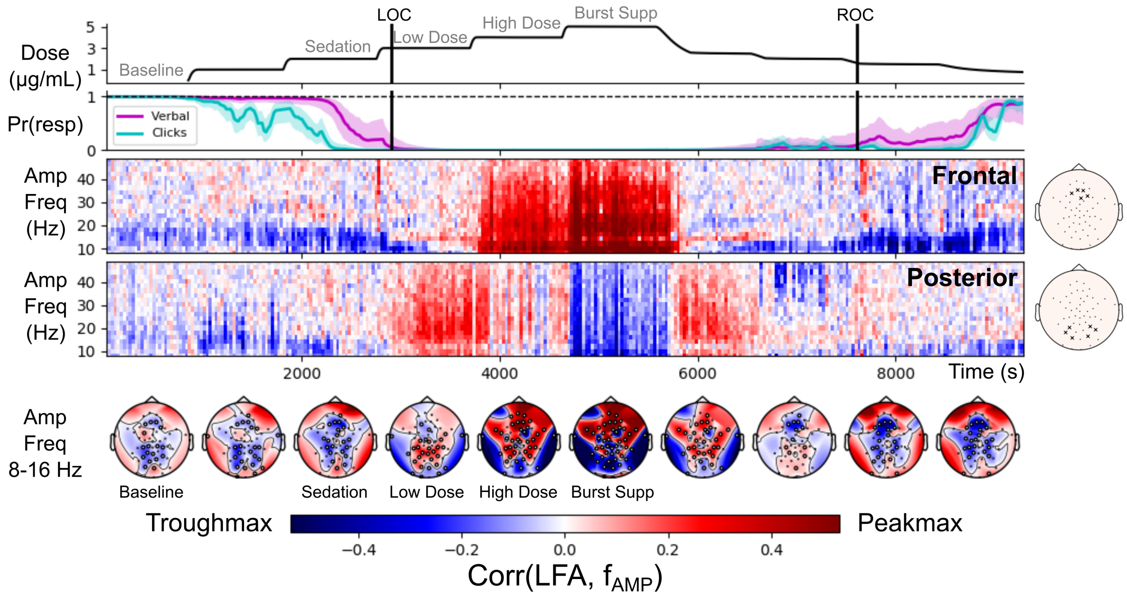


**Fig. S4:** Sensor-level summary for Subject 4. Layout as in Figure 1. Note that this subject entered burst suppression (Burst Supp) at the highest dose of propofol, so the previous level was used for the Unconscious High Dose condition.

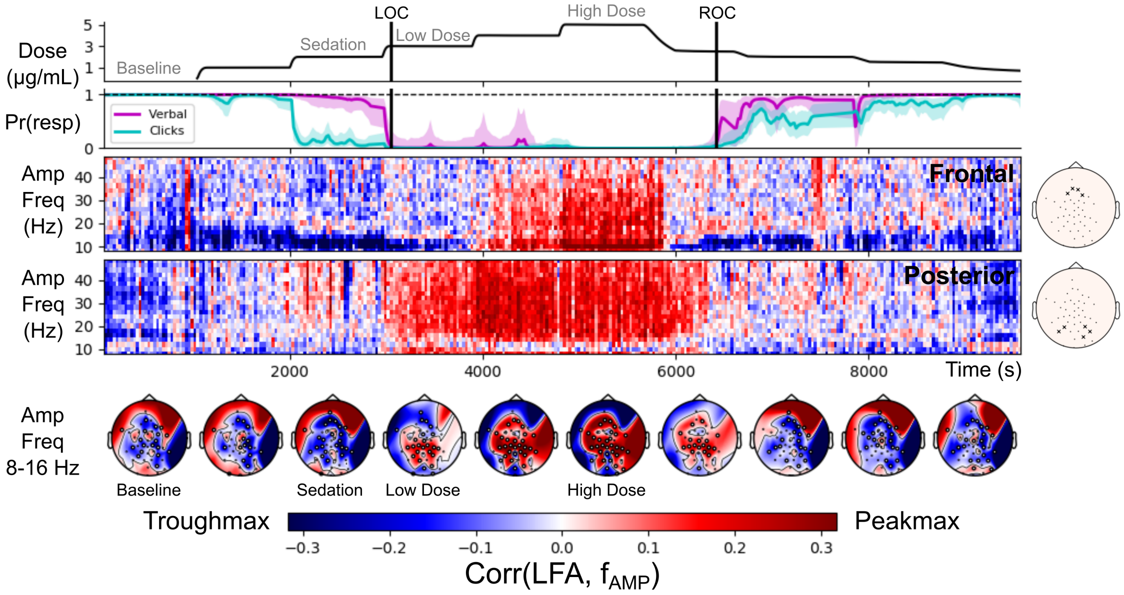


**Fig. S5:** Sensor-level summary for Subject 5. Layout as in Figure 1.

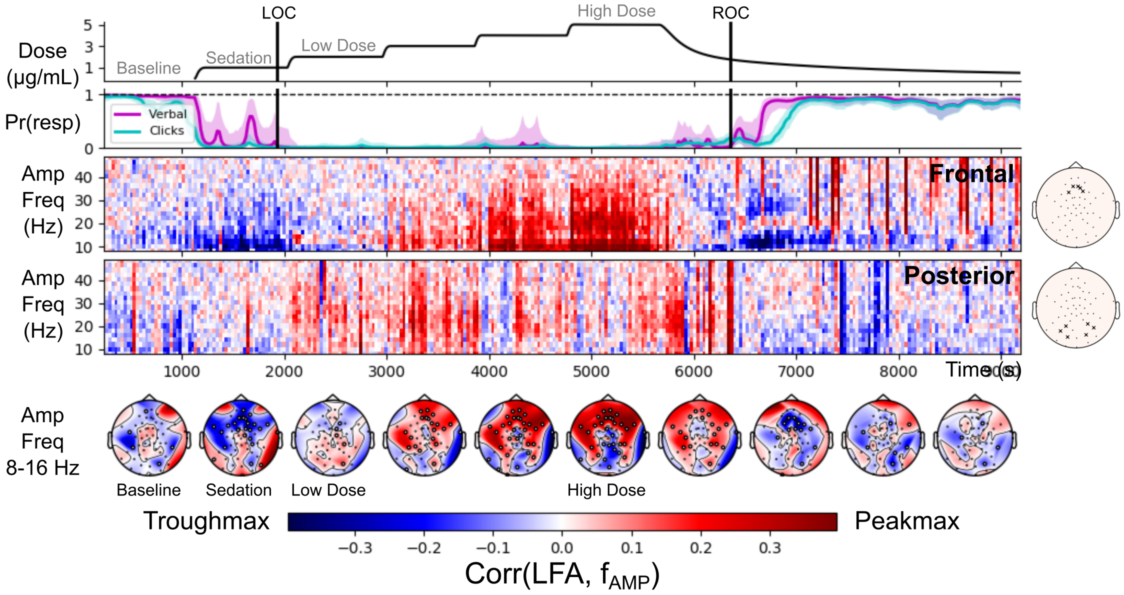


**Fig. S6:** Sensor-level summary for Subject 8. Layout as in Figure 1.

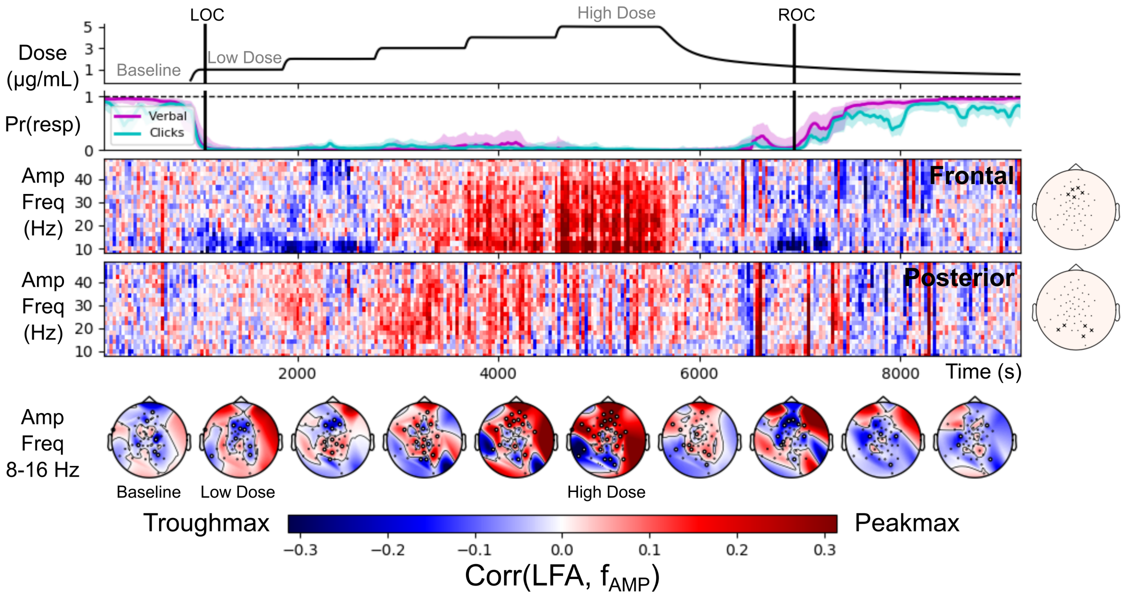


**Fig. S7:** Sensor-level summary for Subject 9. Layout as in Figure 1.

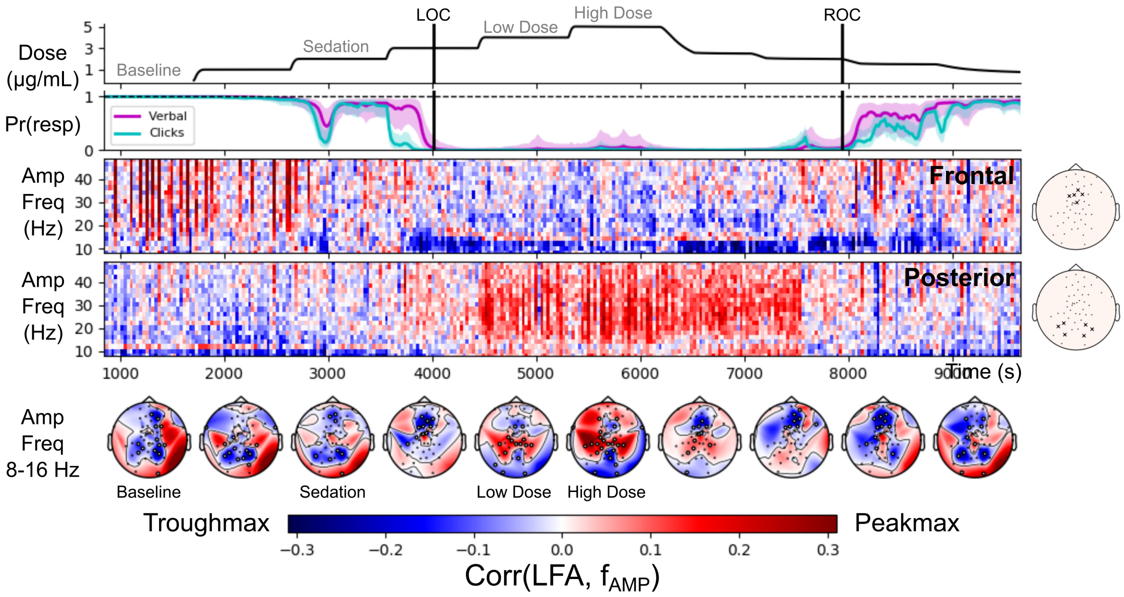


**Fig. S8:** Sensor-level summary for Subject 10. Layout as in Figure 1. Note that the Unconscious High Dose condition for this subject does not exhibit broadband peakmax dynamics over frontal electrodes: we interpret this to indicate that the subject was in a lighter state of unconsciousness than other subjects at the highest propofol dose.

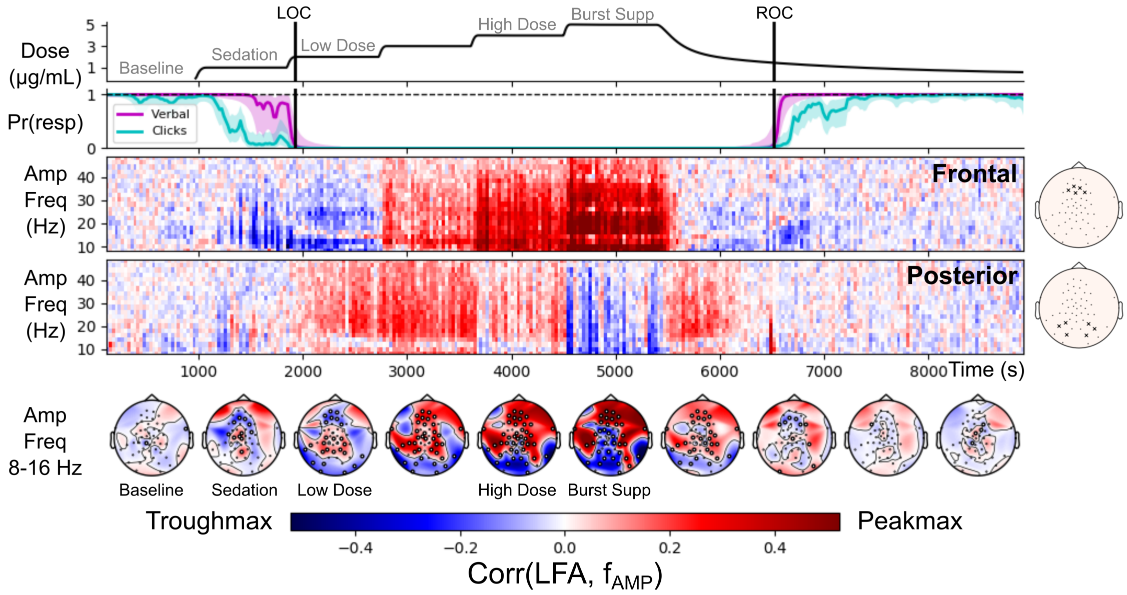


**Fig. S9:** Sensor-level summary for Subject 13. Layout as in Figure 1. Note that this subject entered burst suppression (Burst Supp) at the highest dose of propofol, so the previous level was used for the Unconscious High Dose condition.

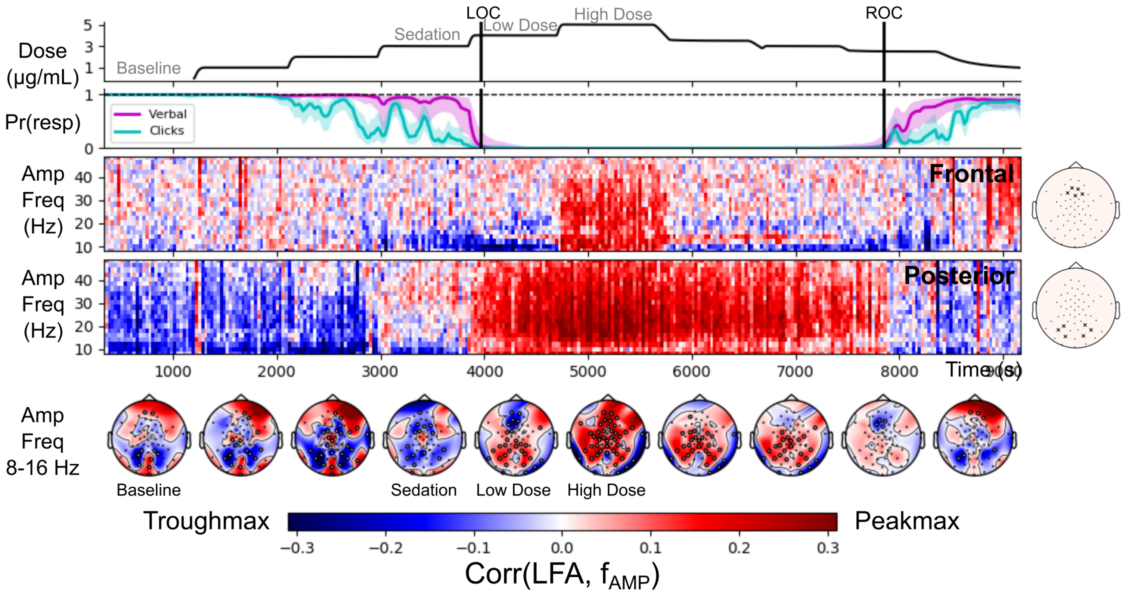


**Fig. S10:** Sensor-level summary for Subject 15. Layout as in Figure 1.

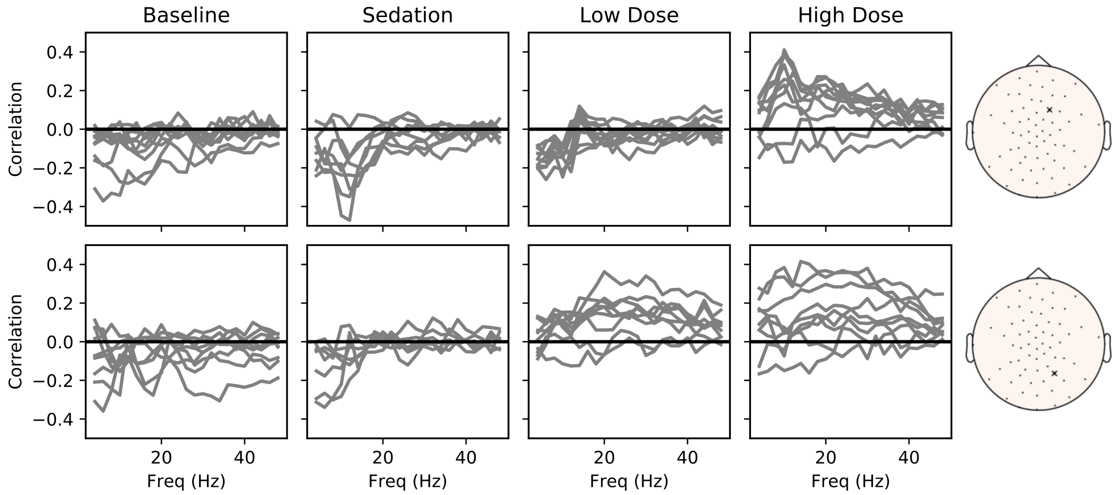


**Fig. S11:** Cross-frequency coupling results for two electrodes (indicated in the insets, right), across all subjects (traces), for the four levels of interest.

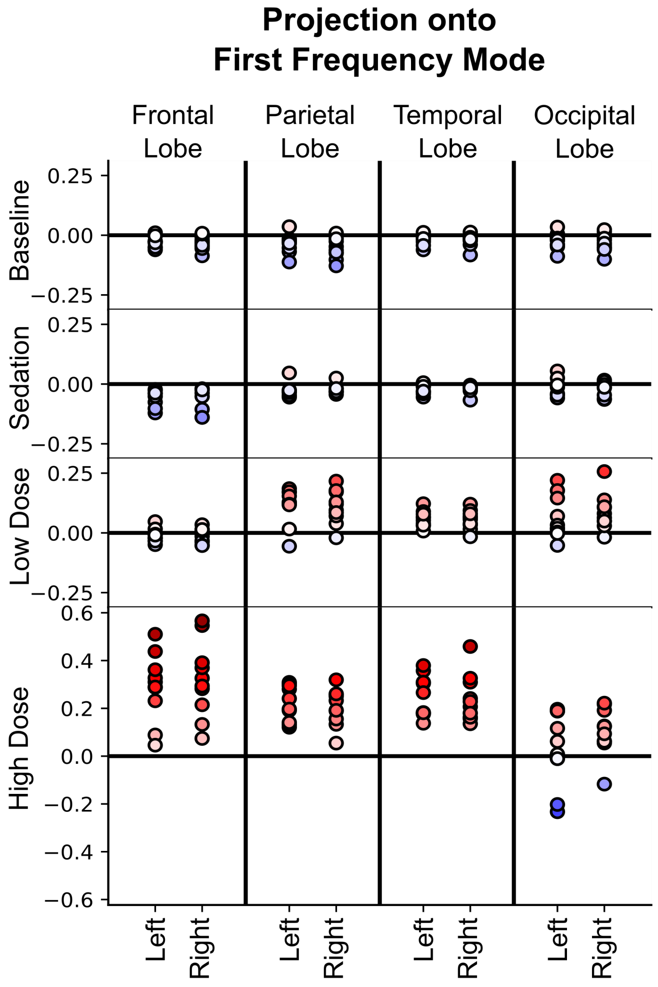


**Fig. S12:** Cross-frequency coupling patterns estimated by lobe in source space, projected onto the first principal mode. Each marker represents one subject. The markers are colored based on the value of the projection (also shown on the y-axis). Because the first mode is positive for all frequencies, positive values (red) correspond to broadband peakmax and negative values (blue) correspond to broadband troughmax.

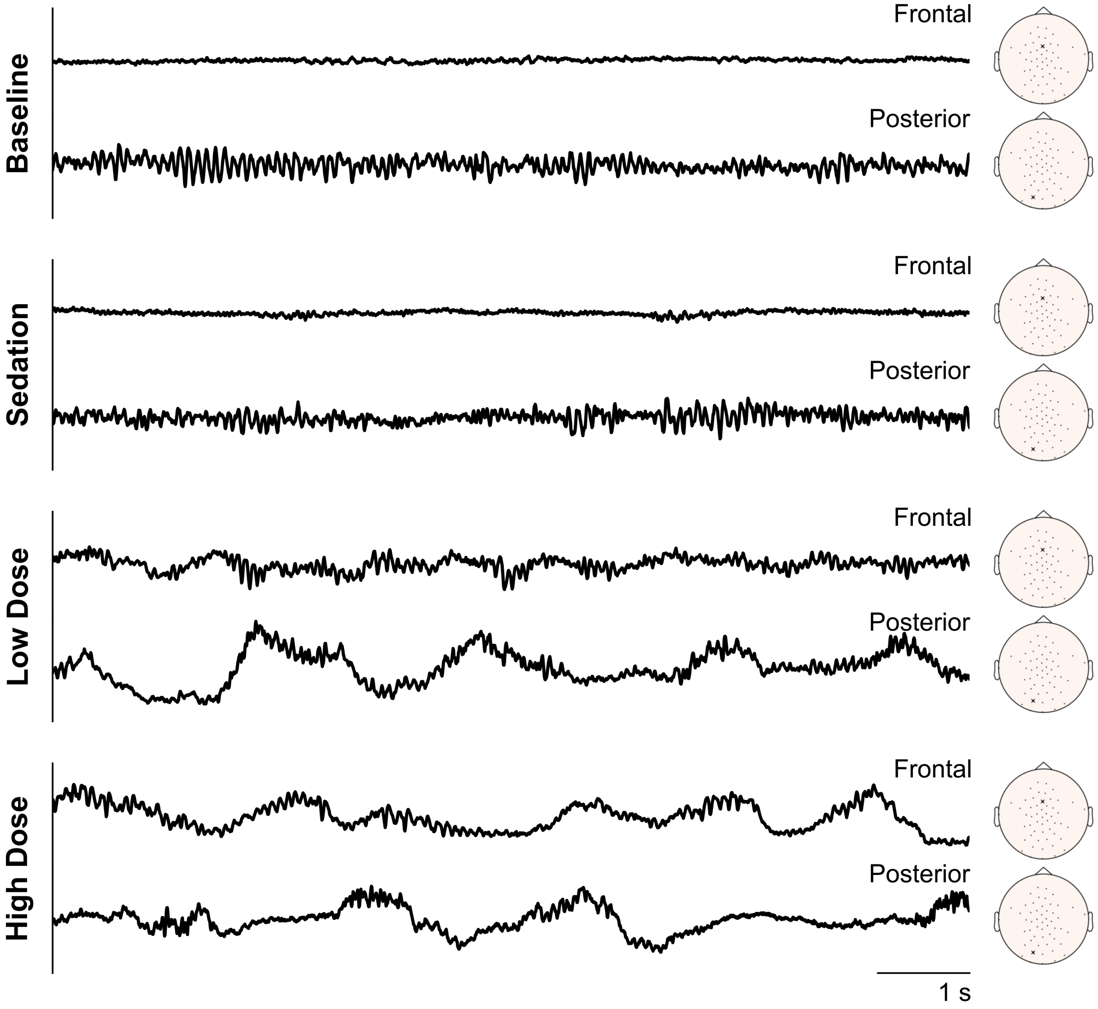


**Fig. S13** Example EEG traces from two electrodes of Subject 7 (see Figure 1). Ten seconds of EEG data were selected from the four levels of interest (Baseline, Sedation, Unconscious Low Dose, and Unconscious High Dose), and the EEG data are displayed for a frontal and a posterior electrode (electrode locations shown in the insets to right). While Baseline and Sedation have little coupling of activity to the slow-wave, Unconscious Low Dose and Unconscious High Dose reflect the cross-frequency coupling evident in Figure 1: at Low Dose, the posterior electrode shows broadband coupling to the peak of the slow-wave; at High Dose, both posterior and frontal electrodes show broadband coupling to the peak of the slow-wave.
